# Ultraviolet-C mediated inactivation of *Candida auris*, a rapid emerging health threat

**DOI:** 10.1101/2023.04.21.537870

**Authors:** Carolina Koutras, Richard L. Wade

**Affiliations:** Department of Clinical Research, R-Zero Systems, Salt Lake City, UT

**Keywords:** Ultraviolet-C, Infection Prevention, *Candida auris*, Disinfection, Patient Safety

## Abstract

Healthcare-associated infections (HAIs), particularly those caused by multidrug-resistant organisms (MDROs), pose significant challenges to patient safety. *Candida auris* (*C. auris*), an emerging MDRO fungus, has been acknowledged as an urgent threat by the Centers for Disease Control and Prevention (CDC) due to its high mortality and difficulty in prevention, diagnosis, and treatment. In this study, we investigated the efficacy of 254 nm ultraviolet-C light (UV-C) in inactivating *C. auris* on hard surfaces. A mobile UV-C tower equipped with high-performance bulbs was used, and within 7 minutes of continuous exposure, ≥ 99.97% (≥ 3.86 log10) inactivation of C. auris was achieved in a patient-room sized test chamber. Our findings suggest that UV-C can serve as an effective infection control measure for preventing *C. auris* and other MDRO HAIs in healthcare settings. Implementation of UV-C disinfection protocols can contribute to enhanced patient safety and combat the growing threat of MDRO pathogens.

## Background

Healthcare associated infections (HAIs) are infectious diseases not present or incubating at admission and contracted within a healthcare environment. They are caused by viral, bacterial, and fungal pathogens, and transmission usually occurs via medical interventions, hospital equipment, and healthcare worker and patient exposure. The most common sites of infection are the bloodstream (central line-associated bloodstream infections or CLABSI), lungs (ventilator-associated pneumonia or VAP), urinary tract (catheter-associated urinary tract infections or CAUTI), and surgical wounds (surgical site infections or SSI). While any microorganism can potentially cause HAIs, there is a growing concern over the high prevalence of cases associated with multidrug-resistant organisms (MDROs) [1]. This increase is due to the misuse and overuse of antibiotics as well as poor hygiene measures and compliance. Commonly seen HAI-causing pathogens include methicillin-resistant *Staphylococcus aureus* (MRSA), vancomycin-resistant *enterococci* (VRE), and *Clostridioides difficile* (C. difficile), but emerging pathogens are also of concern. *Candida auris* (*C. auris*) is an MDRO fungus that can lead to invasive infections and is associated with high mortality. In 2022, the World Health Organization (WHO) included *C. auris* in the top 4 most critical pathogens among 19 of the most concerning fungal pathogens in public health [2]. *C. auris* can only be identified in highly specialized laboratories and is difficult to eliminate and treat. Patients often remain colonized with this potentially deadly pathogen for extended periods of time, sometimes indefinitely, while shedding the fungus and contaminating their immediate environment. Since *C. auris* was first reported in 2016 in the United States, there were 3,270 clinical cases (in which infection is present) and 7,413 colonization cases where individuals carry the organism somewhere on their bodies without signs of active infection. Clinical cases have increased each year, with the most rapid rise occurring during 2020-2021. COVID-19 is believed to have contributed to these numbers, as some patients required intubation and other invasive procedures that put them at higher risk of infection. This month, the Centers for Disease Control and Prevention (CDC) acknowledged *C. auris* as an urgent threat that is difficult, sometimes impossible, to prevent, diagnose and treat [3]. Thus, preventing *C. auris* infections is a key aspect of patient safety in healthcare. Here we investigate the effectiveness of an ultraviolet-C light (UV-C) intervention in reducing *C. auris* on hard surfaces, a common vector of *C. auris* transmission. In under 7 minutes of continuous exposure, UV-C disinfection resulted in > 99.97% inactivation of *C. auris* in a patient-room sized test chamber, suggesting that UV-C can be an effective infection control measure to protect patients from contracting *C. auris* in healthcare.

## Methods

A total of 12 carriers (2.5×5.0 cm glass slides) were inoculated with 0.020 mL of the test culture (*C. auris* CDC AR Bank #0385) in 5% Fetal Bovine Serum (FBS) soil load and dried in a biosafety cabinet at 25.6-25.9°C in a relative humidity of 28%. Three 78” tall UV-C devices (test devices A, B, and C) equipped with 8 high-output lamps, 8 coated posterior parabolic reflectors, 4 long-range passive infrared safety sensors, and delivering approximately 250 mJ/cm2 during a 7-minute cycle at 8 ft (Arc, R-Zero Systems) were placed at 8 feet (∼200 sq feet radius) from the test carriers in a biosafety test chamber. Carriers were exposed to UV-C light for 6 minutes and 59 seconds (+/-1 second) in total; each of the three UV-C devices was tested against their corresponding triplicate test carriers separately. Following harvest of test carriers and three non-exposed control carriers, microbial suspensions were serially diluted in sterile Phosphate Buffered Saline (PBS), plated onto agar growth media using spread plating technique, and quantified as colony forming units (CFUs) at a range of 25-250 CFU/plate. The log10 and percent reductions CFUs were calculated for UV-C exposed carriers relative to controls. The study diagram is shown in Figure 1.

**Figure 1.**
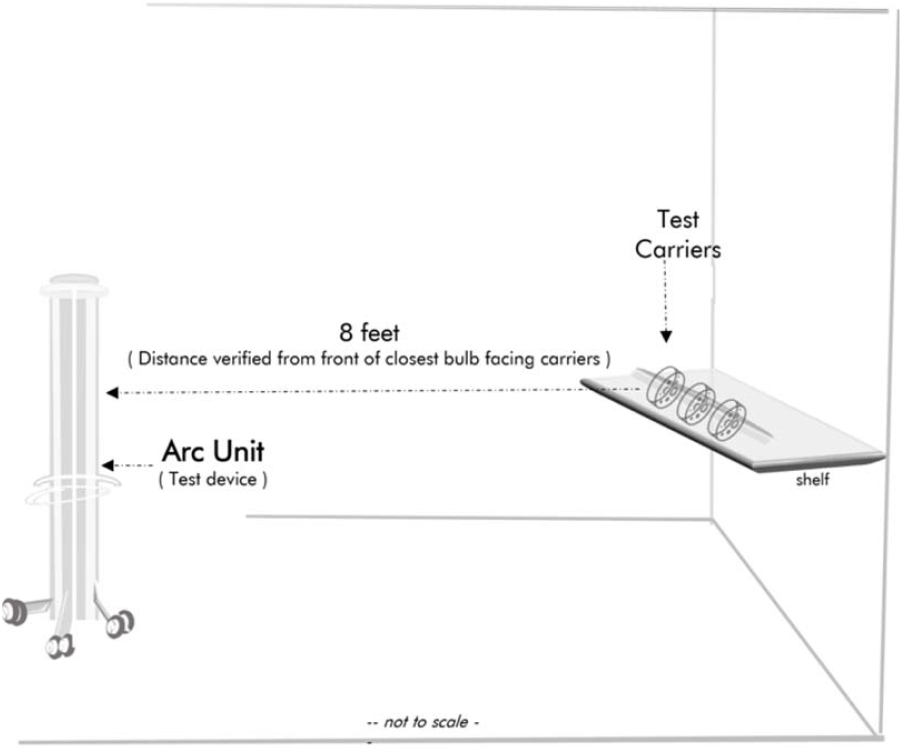
Illustrative Study Diagram. The test device was placed at an 8-feet distance from the test carriers.

## Results

Table 1 summarizes the CFU count per carrier for the controls. Controls had an average CFU count of 1.77E+05 per carrier. Table 2 summarizes the CFU count per carrier following UV-C treatment. Treated carriers averaged a >3.86 log10 reduction and >99.97% percent reduction relative to time zero controls. The lower limit of detection for this study was 1.00E+01 CFU/carrier. Values observed less than the limit are reported as “<1.00E+01” in the results tables. No microbial contamination of any media or test culture was observed during the course of the tests, which were carried out strictly in accordance with Good laboratory Practices (GLP).

**Table 1.**
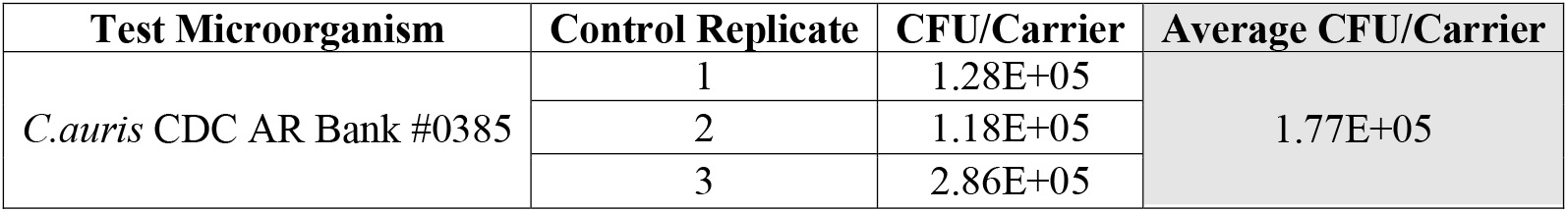
The average CFU count per carrier is shown for the control replicates, not exposed to UV-C light.

| Test Microorganism | Control Replicate | CFU/Carrier | Average CFU/Carrier |
| --- | --- | --- | --- |
| *C.auris* CDC AR Bank #0385 | 1 | 1.28E+05 | 1.77E+05 |
|  | 2 | 1.18E+05 |  |
|  | 3 | 2.86E+05 |  |

**Table 2.**
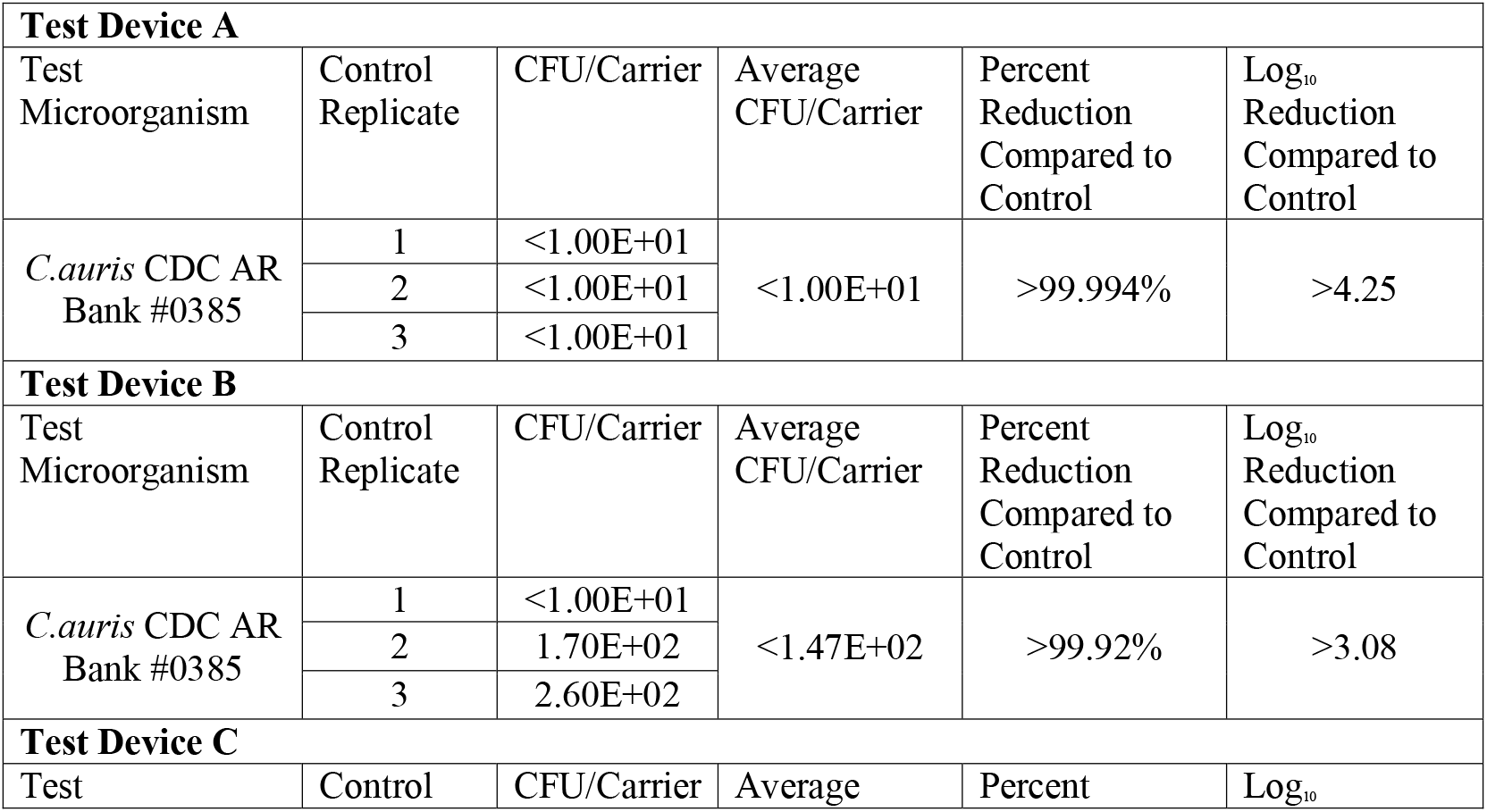
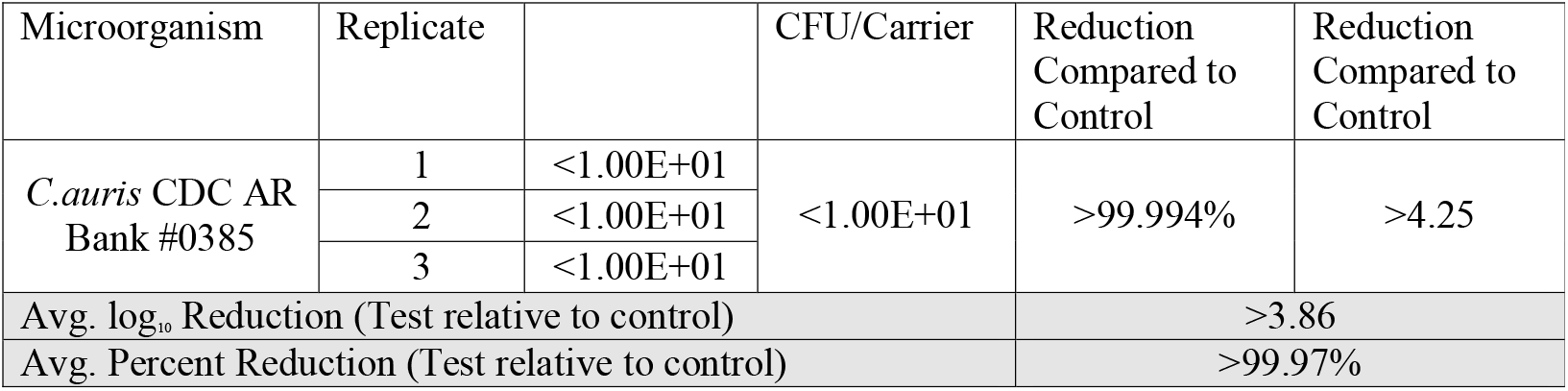
The average CFU count per carrier is shown for the test replicates, exposed to UV-C light.

| Test Device A |  |  |  |  |  |
| --- | --- | --- | --- | --- | --- |
| Test Microorganism | Control Replicate | CFU/Carrier | Average CFU/Carrier | Percent Reduction Compared to Control | Log<sub>10</sub> Reduction Compared to Control |
| *C.auris* CDC AR Bank #0385 | 1 | <1.00E+01 | <1.00E+01 | >99.994% | >4.25 |
|  | 2 | <1.00E+01 |  |  |  |
|  | 3 | <1.00E+01 |  |  |  |
| Test Device B |  |  |  |  |  |
| Test Microorganism | Control Replicate | CFU/Carrier | Average CFU/Carrier | Percent Reduction Compared to Control | Log<sub>10</sub> Reduction Compared to Control |
| *C.auris* CDC AR Bank #0385 | 1 | <1.00E+01 | <1.47E+02 | >99.92% | >3.08 |
|  | 2 | 1.70E+02 |  |  |  |
|  | 3 | 2.60E+02 |  |  |  |
| Test Device C |  |  |  |  |  |
| Test | Control | CFU/Carrier | Average | Percent | Log<sub>10</sub> |

| Microorganism | Replicate |  | CFU/Carrier | Reduction Compared to Control | Reduction Compared to Control |
| --- | --- | --- | --- | --- | --- |
| *C.auris* CDC AR Bank #0385 | 1 | <1.00E+01 | <1.00E+01 | >99.994% | >4.25 |
|  | 2 | <1.00E+01 |  |  |  |
|  | 3 | <1.00E+01 |  |  |  |
| Avg. log<sub>10</sub> Reduction (Test relative to control) |  |  |  | >3.86 |  |
| Avg. Percent Reduction (Test relative to control) |  |  |  | >99.97% |  |

## Discussion

*C. auris*, one of the world’s most feared emerging MDRO pathogens, can lead to invasive fungal disease. It is known to survive on healthcare environmental surfaces for days/weeks and can cause outbreaks [4,5]. In this study, a mobile UV-C tower equipped with high-performance bulbs and reflectors successfully inactivated > 99.97 % (> 3.86 log10) of *C. auris* in less than 7 minutes. Although there are published reports exploring the susceptibility of *C. auris* to UV-C light, significant higher exposure times or shorter exposure distances were required to achieve comparable efficacy as reported in our study [6,7,8]. This highlights the importance of UV-C system design, such as incorporation of bulbs emitting near the peak germicidal wavelength of 254 nm, high UV-C output, and design-optimized UV-C reflectors. Furthermore, even if longer UV-C cycles using lower output bulbs can deliver the same UV-C dose, disinfection protocols incorporating long UV-C interventions are difficult to implement in healthcare. In the largest clinical trial of enhanced UV-C disinfection in healthcare to date, time constraints were identified as a key barrier to implementing enhanced disinfection protocols using UV-C [9]. While this study focused on an emerging pathogen, the germicidal properties of UV-C light are not species-specific. Rather, UV-C inactivates microorganisms by causing damage to their nucleic acids. In conclusion, UV-C disinfection is an effective disinfection technology that can be easily deployed in healthcare for the prevention and control of MDRO pathogens, including *C. auris*.

## Acknowledgements

The authors would like to thank Dr. Benjamin Tanner and Microchem Laboratories (Austin, TX) for their commitment to this project.

